# Altered Thalamocortical Signaling in a Mouse Model of Parkinson’s Disease

**DOI:** 10.1101/2020.07.27.223222

**Authors:** Olivia K. Swanson, David Richard, Arianna Maffei

## Abstract

Activation of the primary motor cortex (M1) is important for the execution of skilled movements and motor learning, and its dysfunction contributes to the pathophysiology of Parkinson’s disease (PD). A well accepted idea in PD research, albeit not tested experimentally, is that loss of midbrain dopamine leads to decreased activation of M1 by the motor thalamus (Mthal). Here, we report that midbrain dopamine loss reduced Mthal input in a laminar- and cell type-specific fashion and induced laminar-specific changes in intracortical synaptic transmission. As a result, M1 activation by Mthal was decreased, but M1 output was increased. Our results demonstrate that loss of midbrain dopaminergic neurons alters thalamocortical activation of M1, and provide novel insights into circuit mechanisms for motor cortex dysfunction in a mouse model of PD.

## Introduction

Skilled voluntary movement is essential for nearly all behaviors. It requires coordinated communication across the motor pathway, which includes the basal ganglia, cerebellum, thalamus and cortex, to produce the intended movement. The primary motor cortex (M1) is the output center of the motor pathway. M1 directly controls movement via the corticospinal-projecting neurons in deep layers (Lemon, 1993), while cells in superficial layers provide feedback to other cortical regions and the striatum (Oswald et al., 2013; Shepherd, 2013). The activity of M1 pyramidal neurons is regulated by neighboring inhibitory neurons, including parvalbumin-expressing (PV^+^) cells. Engagement of M1 excitatory and inhibitory cells is required for many aspects of motor function, including movement execution (Kaufman et al., 2013; Melzer et al., 2017), motor planning (Svoboda and Li, 2018) and learning (Hosp et al., 2011; Chen et al., 2015; Biane et al., 2016; Kida et al., 2016).

A major long-range input to M1 arises from the ventroanterior/ventrolateral nuclei of the thalamus (Mthal). Mthal projections to M1 are most dense in L2/3 and L5 (Hooks et al., 2013; Hunnicutt et al., 2014) and make glutamatergic synapses with excitatory and inhibitory neurons (Biane et al., 2016; Shigematsu et al., 2016). The synaptic properties of this input are not as well understood as thalamic projections to sensory cortices. Findings from sensory cortices suggest that driver pathways such as Mthal-M1 to elicit large, depressing postsynaptic currents in both excitatory and inhibitory cells (Swanson and Maffei, 2019). Feedforward inhibition in particular is not well studied in M1, although in other sensory cortices it plays a central role in cortical processing (Gabernet et al., 2005; Wang et al., 2010). In M1, it may play a crucial role in gating cortical activation and ultimately movement.

Transient disruption of M1 activity diminishes control over voluntary movement (Schieber and Poliakov, 1998; Stepniewska et al., 2014; Chen et al., 2019), and chronic changes in M1 activity have been associated with movement disorders, including Parkinson’s disease (PD). PD is a highly prevalent movement disorder characterized by progressive tremor, rigidity, akinesia, and postural instability (Wenning et al., 2005; Ostrem and Galifianakis, 2010; Hirsch et al., 2016). While it is well-established that the loss of dopaminergic neurons in the substantia nigra pars compacta (SNc) leads to dramatic shifts in synaptic transmission in many motor areas (Day et al., 2006; Bagetta et al., 2010; Fan et al., 2012), the effects on M1 remains understudied. Emerging reports show evidence of abnormal neural activity within M1 in PD (Lindenbach and Bishop, 2013; Calabresi and Di Filippo, 2015). A working hypothesis in the field is that loss of midbrain dopamine impacts thalamocortical excitation of M1, yet the synaptic underpinnings of this dysfunction remain unclear.

Here, we performed whole-cell recordings of M1 pyramidal neurons and inhibitory PV^+^ interneurons to determine how loss of midbrain dopaminergic neurons impacts Mthal synaptic transmission. We used optogenetic stimulation of the Mthal-M1 pathway in a mouse model of PD and measured synaptic drive and dynamics in a dopamine-depleted state. Our results indicate that midbrain dopamine depletion leads to layer-specific reduction of Mthal input to M1 pyramidal neurons, while Mthal activation of the inhibitory circuit mediated by PV^+^ cells is preserved. Additional shifts in excitation/inhibition balance lead to increased pyramidal neurons output in the face of reduced Mthal drive. These are the first direct experimental evidence that Mthal transmission is reduced due to loss of dopaminergic input to the motor pathway, and provide a mechanistic framework for M1 dysfunction in PD.

## Methods

### Animals

All experimental procedures were designed and executed following the guidelines of the National Institute of Health and were approved by Stony Brook University’s Institutional Animal Care and Use Committee. Mice of both sexes were used in all experiments. Most recordings of excitatory neurons were collected from C57BL/6NCrl (#027, Charles River) animals. Recordings of fast-spiking interneurons, as well as a subset of pyramidal neurons, were made from acute slice preparations obtained from the progeny (Pvcretdtomato) of female PV-Cre (B6;129P2-*Pvalb^tm1(cre)Arbr^*/J, #00809, Jackson Laboratories) and male Ai14 (B6.Cg-*Gt(ROSA)26Sor^tm14(CAG-tdTomato)Hze^*/J, #007914, Jackson Laboratories) mice. The number of animals and number of cells within each experimental group are reported as *N* and *n,* respectively.

### Surgical Procedures

Animals at P36-P50 were anesthetized with a ketamine/xylazine cocktail prepared in sterile saline (100mg/kg ketamine, 10mg/kg xylazine), and received an IP injection of desipramine (1.25mg/mL, 20mL/kg) 30 minutes prior to surgery. Each animal received a unilateral injection of 6-hydroxydopamine hydrochloride (6OHDA) centered on the SNc and an injection of AAV9-CAG-ChR2-Venus (diluted to final titre 1.127×10^12^ GC/mL, Penn Vector Core; ChR2 (Petreanu et al., 2007)) in the VA/VL of the same hemisphere. Following the opening of craniotomies over each injection site, a pressure injection system (Nanoinject, Drummond) was used to inject 50nL of virus in into the VA/VL (Bregma-0.7, midline ± 1.0, surface-3.75) at 4.6nL intervals. The injection pipette was left in place for 5min to prevent virus spread up the needle tract. 6OHDA was prepared fresh (15mg/mL in 0.02% ascorbate saline) and injected at each of two injection sites (250nL, 2.75μg at each site) centered along the anterior-posterior extent of the SNc (Bregma+1.2/0.8, midline-1.1, surface-0.8). Control animals were injected with equivalent volumes of drug vehicle. Following surgery, animals recovered on a heating pad and were monitored daily for food and water intake.

### Cylinder motor task

Prior to preparing slices, animals were assessed for motor impairment via a cylinder motor task (Iancu et al., 2005). Each animal was placed in a clear acrylic cylinder surrounded by mirrors, with a camera positioned overhead, and allowed to freely explore for 10min. Exploratory mouse movements were filmed and analyzed by an experimenter blind to experimental conditions. Forelimb impairment was expressed as the ratio of weight-bearing wall touches made by the limb contralateral to the SNc injection and total wall touches made by both forelimbs.

### Slice electrophysiology

2-3 weeks after surgery, animals were anesthetized with isoflurane using the bell jar method and rapidly decapitated. The brain was dissected and sectioned in ice cold oxygenated ACSF using a vibrating blade microtome (Leica VT1000S). 300μm slices containing forelimb M1 (Tennant et al., 2011) were transferred into 37°C ACSF to recover for 30min, then moved into room temperature ACSF to stabilize for 1hr. Whole-cell patch clamp was performed at room temperature and guided by DIC optics. Recordings were obtained with pulled borosilicate glass pipettes with a resistance of 3-4MΩ, filled with internal solution containing the following (in mM): 100 K-Glu, 10 K-HEPES, 4 Mg-ATP, 0.3 Na-GTP, 10 Na-phosphocreatine, and 0.4% biocytin, pH 7.35 titrated with KOH and adjusted to 295 mOsm with sucrose (E_rev_[Cl^−^] = −49.8mV), unless otherwise noted. PV^+^ interneurons were targeted under fluorescent light in Pvcretdtomato animals. The firing pattern of each cell was assessed in current clamp by somatic current injection (700ms duration, −100 to 450pA in 50pA steps). Mthal terminal fields were stimulated with 1ms pulses of blue light (470nm), produced by a high-powered LED lamp (X-cite 120LEDMini, Excitalas), passed through a filter and delivered through a 40x water immersion objective. Intensity of ChR2 stimulation was controlled by the LED dial, and the intensity of light emitted from the objective was determined using a power meter (PM100D, Thorlabs). Light pulses were delivered at 1%, 5%, 10%, 15%, and 20% LED intensity (in mW emitted from objective: 0.47, 1.04, 1.61, 2.20, 2.78, respectively). To assess input/output curves, stimulation was delivered for 5 sweeps (30s between each sweep, at each intensity), while excitatory postsynaptic currents (EPSC) were recorded in voltage clamp. Short-term dynamics of Mthal input were assessed with optical stimulation at 5Hz and 10Hz at 5% LED intensity, and responses were recorded in voltage and current clamp. We measured spontaneous synaptic currents onto M1 neurons by recording neurons in voltage clamp with an internal solution containing (in mM): 20 KCl, 100 Cs-sulfate, 10 K-HEPES, 4 Mg-ATP, 0.3 Na-GTP, 10 Na-phosphocreatine, 3 QX-314 (Tocris Bioscience), 0.2% biocytin (E_rev_[Cl^−^] = −49.8mV)). Isolated spontaneous excitatory currents (sEPSC) were measured while holding cells at the reversal potential for chloride (−50mV), while isolated spontaneous inhibitory currents (sIPSC) were measured at the reversal potential for excitatory cations (+10mV). Artificial Cerebrospinal Fluid (ACSF) used in all electrophysiology experiments contained the following (in mM): 126 NaCl, 3 KCl, 25 NaHCO_3_, 1 NaHPO_4_, 2 MgSO_4_, 2 CaCl_2_, and 14 dextrose. Exctiability measurements (input resistance, frequency-current curves), were calculated from current clamp recordings where cells received somatic current injections of increasing intensity (− 100pA to 450pA, 50pA steps). Series resistance (R_s_) was tracked throughout the experiment and data from cells whose R_s_ was >10% of their input resistance or changed >20% over the course of the experiment were excluded.

### Immunohistochemistry

Tissue containing injection sites, and all slices used for recordings were post-fixed in 4% paraformaldehyde (0.01M phosphate buffered saline (PBS), pH 7.4) for one week. Brain tissue containing injection sites were sectioned in the coronal plane at 50μm thickness with a vibrating blade microtome (Leica VT1000S). Tissue sections processed with immunofluorescent protocols were rinsed in PBS 3 times, for 10min each (3×10min), then incubated in an antigen retrieval solution (10mM sodium citrate in dH2O, pH 8.5, at 45°C, 30min for 50μm sections and 45min for 300μm sections). Tissue was rinsed again in PBS 3×10min, then incubated in glycine (50mM, in PBS, 1hr for 50μm sections, 2hr for 300μm sections). Following additional 3×10min PBS rinses, the tissue was incubated in a pre-block solution (in PBS: 5% bovine serum albumin (Sigma), 5% normal goat serum (Vector Laboratories), 0.2%/1% triton-x (VWR) for 50μm/300μm sections) for 1hr or 3hrs at room temperature for 50μm or 300μm sections, respectively. Following pre-block, sections were incubated overnight at 4°C in an antibody incubation solution (in PBS: 1% bovine serum albumin (Sigma), 1% normal goat serum (Vector), 0.1% triton-x (Sigma)) containing the appropriate primary antibodies and streptavidin reagents (Table 1A-C). The following day, tissue was rinsed in 3×10min in PBS and transferred into the antibody incubation solution containing secondary antibodies (Table 1A-C). 300μm slices were incubated in secondary antibodies for 6hr at room temperature, and 50μm slices were incubated for 4hr. Following a 3×10min PBS rinse, 300μm slices were counterstained with Hoechst33342 (1:5000, Invitrogen H3570) for 20min and 50μm sections containing ChR2 injection sites were counterstained with Neurotrace (435/455) for 30min, then rinsed in 0.1M phosphate buffer (PB), mounted, and coverslipped with fluorescent mounting medium (Fluoromount-G, ThermoFisher).

**Table 1.** Details for antibodies used in immunohistochemistry. **(A)** List of primary antibodies, streptavidin, and secondary antibodies used to process 300μm sections used for whole-cell recordings in C57BL/6 mice. **(B)** List of primary antibodies, streptavidin, and secondary antibodies used to process 300μm sections used for whole-cell recordings in PVcretdtomato mice. **(C)** List of primary and secondary antibodies used to process 50μm sections containing ChR2 injection sites in C57BL/6 and PVcretdtomato mice. **(D)** List of primary and secondary antibodies used to process 50μm sections containing 6OHDA/vehicle injection sites in C57BL/6 and PVcretdtomato mice.

| <u>Antibody</u> | <u>Host Species</u> | <u>Dilution</u> | <u>Vendor</u> | <u>Catalog Number</u> |
| --- | --- | --- | --- | --- |
| anti-GAD67 | mouse | 1:500 | Millipore | MAB5406 |
| Streptavidin Alexa Fluor 568 | --- | 1:2000 | ThermoFisher | S11226 |
| anti-GFP | chicken | 1:5000 | Abcam | ab13970 |
| anti-mouse Alexa Fluor 647 | goat | 1:500 | ThermoFisher | A-21235 |
| anti-chicken Alexa Fluor 488 | goat | 1:500 | Abcam | ab150173 |

**B. 300µm recorded slices from PVcretdtomato:**
| <u>Antibody</u> | <u>Host Species</u> | <u>Dilution</u> | <u>Vendor</u> | <u>Catalog Number</u> |
| --- | --- | --- | --- | --- |
| Streptavidin Alexa Fluor 647 | --- | 1:2000 | ThermoFisher | S21374 |
| anti-GFP | chicken | 1:5000 | Abcam | ab13970 |
| anti-chicken Alexa Fluor 488 | goat | 1:500 | Abcam | ab150173 |

**C. 50µm sections for Chr2 Injection Site:**
| <u>Antibody</u> | <u>Host Species</u> | <u>Dilution</u> | <u>Vendor</u> | <u>Catalog Number</u> |
| --- | --- | --- | --- | --- |
| anti-GFP | chicken | 1:5000 | Abcam | ab13970 |
| anti-chicken Alexa Fluor 488 | goat | 1:500 | Abcam | ab150173 |

**Table 1. Details for antibodies used in immunohistochemistry.**
| <u>Antibody</u> | <u>Host Species</u> | <u>Dilution</u> | <u>Vendor</u> | <u>Catalog Number</u> |
| --- | --- | --- | --- | --- |
| anti-Tyrosine Hydroxylase | rabbit | 1:1000 | Abcam | ab112 |
| anti-rabbit, biotinylated | goat | 1:200 | Vector Laboratories | BA-1000 |

Sections through the SNc and VTA, containing 6OHDA injection sites, were processed and visualized in 3’-3’-diaminobenzidine (DAB). This tissue underwent the described protocol, with the following modifications: following glycine incubation, sections were rinsed 3×10min in PBS and then incubated in 0.3% H2O2 (in PBS, 30min incubation at 4°C). Sections were then rinsed 3×5min in PBS, then blocked for endogenous avidin-biotin reactivity (Avidin/Biotin Blocking Kit, Vector Laboratories). Following 3×1min rinses, tissue was placed in the preblocking solution, then primary antibodies (Table 1D). After secondary antibody incubation, sections were rinsed 3×10min in PBS and then incubated in avidin-biotin horseradish peroxidase (Vectastain Elite ABC kit, Vector) for 1hr at room temperature. Following an additional 3×10min rinse, sections were developed for 60s in 3,3’-diaminobenzidine (DAB Peroxidase Substrate Kit, Vector). Sections were rinsed a final time for 3×10min in 0.1M phosphate-buffered saline (pH 7.4), mounted on gelatin-coated slides, and air-dried for one week. Slides were then dehydrated in a series of alcohols (70%, 95%, 100%), cleared in xlyenes, and coverslipped with Entellan mounting medium. Imaging of fluorescently labeled sections was performed on a laser-scanning confocal microscope (Olympus), and brightfield images were obtained using a widefield microscope (Olympus).

### Animal Inclusion Criteria

Sections containing the SNc, as well as more anterior sections containing the striatum, were processed for TH immunoreactivity and imaged. These images were used as inclusion criteria for an animal to remain in the study. Lesioned animals included in these data showed severe dopaminergic cell loss along the extent of the SNc, as well as significant loss of TH^+^ afferents in the striatum. Animals with injection sites too medial, leading to complete loss of VTA dopaminergic neurons, or too posterior, leading to loss of noradrenergic neurons in the locus coeruleus, were excluded from the study. All animals in this study showed significant forelimb motor impairment as tested with the cylinder motor task prior to slice preparation for patch clamp recording. Sections containing thalamus were processed for GFP immunoreactivity to enhance the visualization of ChR2 expression and imaged. ChR2 injection site expression that extended into adjacent thalamic nuclei, particularly the ventroposteromedial nucleus or the posteromedial nucleus, or expression in the needle tract in forelimb motor cortex, were criteria for exclusion from the study.

### Data Analysis

Measurements from collected electrophysiological recordings were performed using custom-made procedures in Igor (Wavemetrics), as well as using a template-matching system in Clampfit (Molecular Devices). For analysis of input/output curves, EPSC amplitude was measured as the absolute difference between the pre-light pulse holding current (averaged across 100ms) and the absolute minima of the EPSC. EPSC coefficient of variation (1/CV^2^) and variance to mean ratio (VMR) were measured using EPSCs evoked at 5% LED intensity, as previously described (van Huijstee and Kessels, 2020). Briefly, 1/CV^2^ was calculated as the squared inverse of the standard deviation of EPSC amplitude, divided by the mean amplitude (averaged across 5 sweeps/cell), for each cell, and VMR was calculated as the variance of the EPSC amplitude (averaged across 5 sweeps/cell) divided by the mean. EPSC charge and decay tau at 5% LED intensity were calculated in Clampfit. Analysis of short-term dynamics in either voltage clamp or current clamp was performed in Igor by measuring the amplitude of each event. For current clamp traces where neurons fired in response to TC stimulation, the number of action potentials occurring in the 100ms window following each light pulse were counted. In cases where the neuron responded with subthreshold excitatory postsynaptic potentials (EPSPs), the amplitude of each EPSP was calculated as the difference between the pre-light pulse membrane potential (averaged across 100ms) and the absolute maxima in the 100ms window following each light pulse. The amplitude and instantaneous frequency of spontaneous EPSCs and IPSCs were measured in Clampfit using custom template matching criteria, and these data were organized into cumulative probability plots in Igor (100 events per cell). Dynamic input resistance was calculated as the slope of the current-voltage (IV) curve obtained from a series of hyperpolarizing current steps (−100pA to 0pA). Frequency-current (*fI*) curves were computed as the average action potential frequency during a given current step across all cells in each experimental group (50pA to 450pA). Following initial data analyses in Igor and Clampfit, data were compiled in Microsoft Excel. For figures where each data point was representing measurements from a single animal rather than a single cell, data collected from neurons located within each layer of the given animal were averaged, and the mean from each animal was then used in any statistical tests performed.

### Statistical Analyses

Statistical tests were performed in Microsoft Excel and the add-in statistical program XLSTAT. All data are shown as mean ± standard error of the mean (SEM) for the number of neurons (*n*) and the number of animals (*N*) indicated. Unpaired 2-tailed Student’s t-tests were used to test for statistically significant differences in mean between experimental groups, and differences between cumulative distributions were assessed with the Kolmogorov-Smirnov test. P-values ≤0.05 were considered significant.

## Results

### Targeting Mthal input onto M1 neurons in a mouse model of PD

Each animal received a unilateral injection of AAV9-CAG-ChR2-Venus into VA/VL (Mthal), as well as two ipsilateral injections of 6OHDA, centered along the anterior-posterior extent of the SNc (**Fig. 1A**). Immunohistochemical enhancement of the injection site and terminal fields revealed targeted ChR2 injections of Mthal and the expected double-banded thalamocortical axon distribution in M1 (**Fig. 1B**) (Hooks et al., 2013; Biane et al., 2016). Recorded M1 neurons were filled with biocytin and stained with fluorescently-conjugated streptavidin to confirm pyramidal neuron morphology and laminar location (**Fig. 1B**). At the time of recording, all lesioned animals showed significant motor impairment, expressed as biased use of the forelimb ipsilateral to the 6OHDA injection (**Fig. 1C**, wall touch ratio: Vehicle = 0.48±0.025, 6OHDA = 0.25±0.039, p=7.8E^−5^). Sections containing midbrain dopaminergic nuclei were stained for tyrosine hydroxylase (TH), and animals with absence of TH^+^ neurons within the SNc of the injected hemisphere were considered successfully lesioned (**Fig. 1D**).

**Figure 1.**
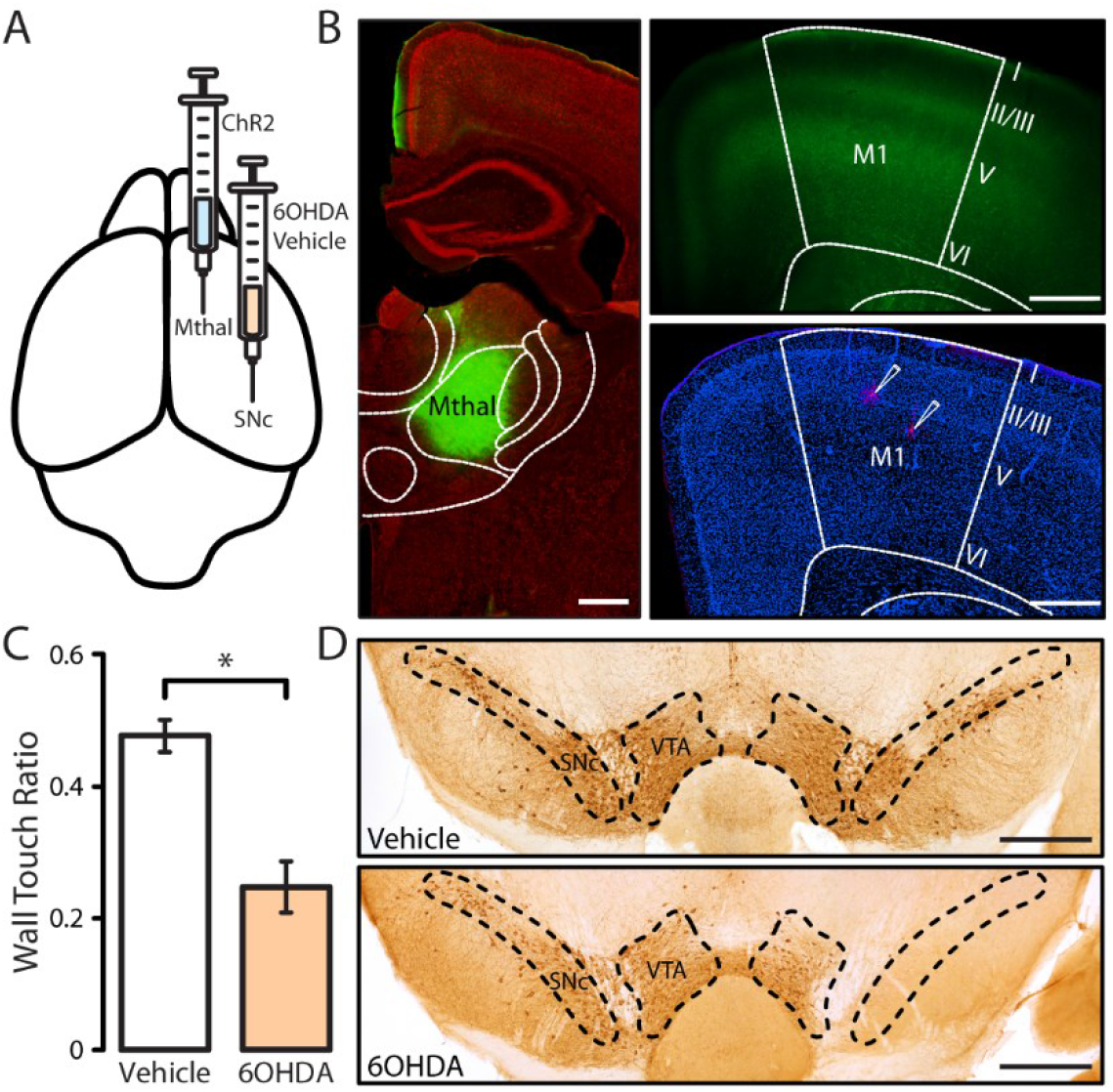
Experimental setup. **(A)** Schematic showing dual, ipsilateral injections of ChR2 into Mthal and 6OHDA/vehicle into the SNc. **(B)** Left panel: 50μm coronal section of thalamus (red: Neurotrace) containing ChR2 injection site (green) within Mthal. Top right panel: 300μm recorded slice with ChR2 (green) expression producing a double-banded terminal field pattern in M1. Bottom right panel: 300μm recorded slice of containing two recorded pyramidal neurons (red: Streptavidin, blue: Hoechst). **(C)** Cylinder motor assessment results. Wall touch ratio calculated as touches with forelimb contralateral to SNc injection divided by total forelimb use. (Vehicle N=17, 6OHDA N=15, *denotes p<0.05). **(D)** Coronal midbrain sections stained for tyrosine hydroxylase immunoreactivity and visualized with DAB. All scale bars = 500μm.

### Reduced thalamocortical drive onto L2/3 pyramidal cells following dopamine depletion

Expression of ChR2 within Mthal produces a double banded terminal field pattern within M1, with the densest ChR2-expressing axons in L2/3 as well as L5. Each layer plays a unique role in the circuit of M1, with L2/3 acting primarily as an input and integration layer while L5 serves more as an output layer (Weiler et al., 2008; Oswald et al., 2013). With their distinct functions in mind, we asked if loss of dopaminergic signaling in the motor pathway may have laminar-specific effects on Mthal input to pyramidal neurons in L2/3 and L5.

First, we examined the effect of dopamine depletion on the Mthal-M1 synapses onto L2/3 neurons. All L2/3 cells included in these experiments exhibited a typical pyramidal cell firing pattern (**Fig. 2A**) and were negative for GAD67 immunoreactivity (**Fig. 2B**). To first examine the baseline parameters of this input, single light pulses (1ms) were delivered at 5% LED intensity and responses were recorded in voltage clamp. Optical activation of Mthal axons elicited excitatory postsynaptic currents (EPSCs) in L2/3 neurons of both vehicle-injected and lesioned animals (**Fig. 2C**). At a 5% LED intensity (see Methods), the evoked EPSC amplitude was significantly reduced in L2/3 neurons of lesioned animals (**Fig. 2C** and second data point in panel H, in pA: Vehicle = 563.92±75.41, 6OHDA = 183.65±34.56, p=0.0002). Further reflecting the reduction in EPSC magnitude, the evoked EPSC charge was significantly reduced in L2/3 neurons in the lesioned group relative to controls (**Fig. 2D**, in pA∙ms: Vehicle = 12415.55±1918.97, 6OHDA = 3986.25±669.38, p=0.0011). The kinetics of the evoked EPSC were also altered: EPSC decay time constant (τ), when normalized to the respective EPSC amplitude, was significantly longer in lesioned animals (**Fig. 2E**, in ms/pA: Vehicle = 0.032±0.0087, 6OHDA = 0.086±0.021, p=0.014). These data suggest depletion of the dopaminergic activity in the motor pathway leads to reduced baseline strength and slower current decay at the Mthal input to M1 L2/3 excitatory neurons.

**Figure 2.**
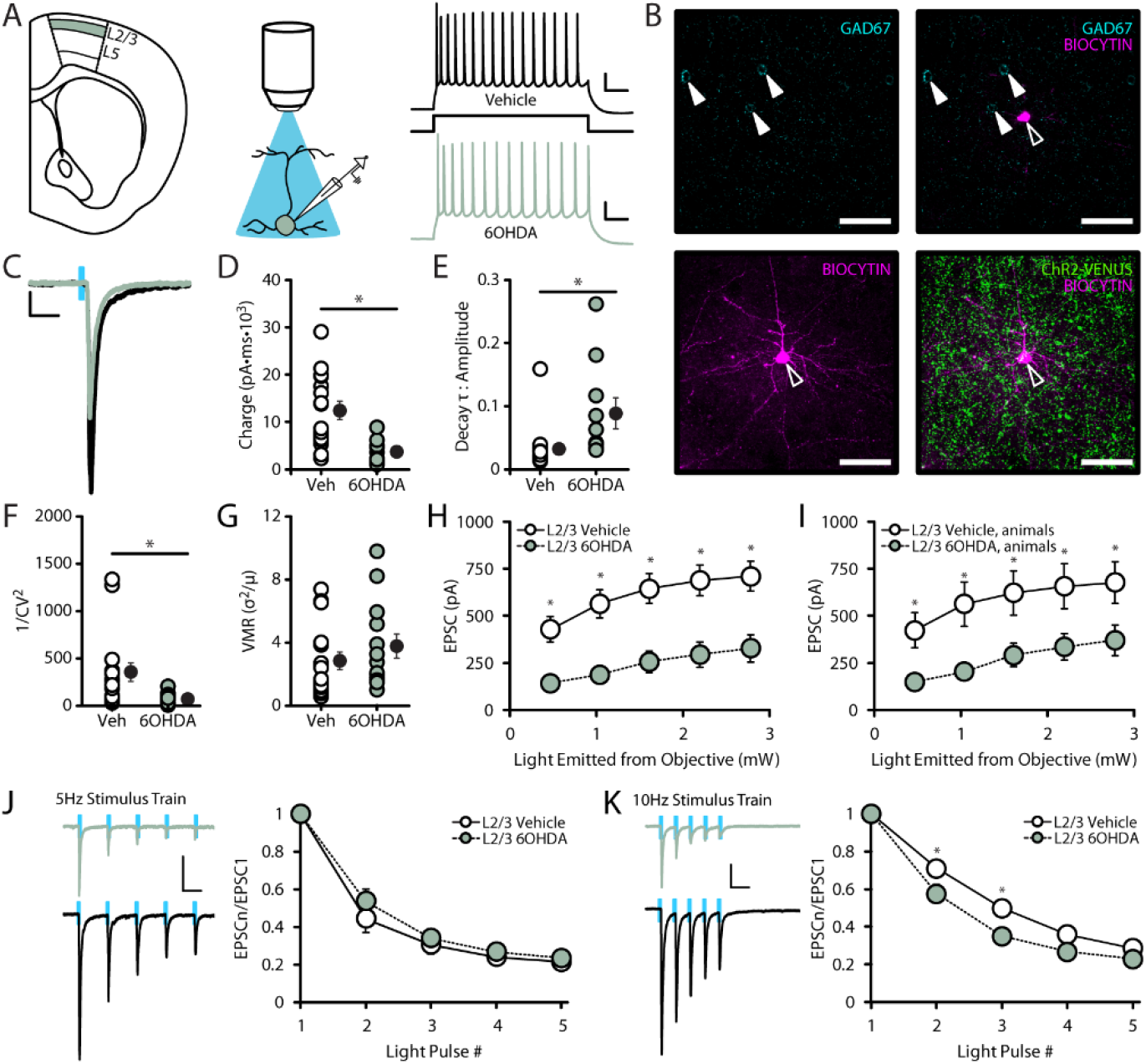
Mthal input to M1 L2/3 neurons is reduced in dopamine-depleted animals. **(A)** Whole-cell recordings of L2/3 pyramidal neurons in M1, with optical activation of ChR2-expressing thalamocortical axons. Right panel: typical firing pattern of a L2/3 neuron in vehicle-injected and lesioned animal. Scale bars = 20V∙100ms **(B)** Confocal images at 60x of an example recorded L2/3 neuron. Top panel: GAD67^−^recorded neuron (open arrow) with adjacent unrecorded GAD67^+^ neurons (filled arrows) and single z-plane depth. Bottom panel left: L2/3 recorded neuron shown as a collapsed stack spanning cell’s full structure. Bottom panel right: merge of L2/3 neuron and surrounding ChR2-expressing axons. Scale bars = 50μm. **(C)** Superimposed example traces of thalamocortical EPSCs from a vehicle-injected (black) and lesioned (green) animal. Scale bar = 50pA∙50ms. **(D)** Calculated EPSC charges, and the group averages (black circles) using 5% LED intensity. **(E)** Calculated EPSC decay tau, normalized to EPSC amplitude, and the group averages (black circles) using 5% LED intensity. **(F)** Average coefficient of variation (1/CV^2^) of EPSCs in response to 5% LED intensity. **(G)** Average variance-to-mean raito (VMR) of EPSCs in response to 5% LED intensity. **(H-I)** Average input-output curves of EPSC amplitude in response to increasing LED intensity. **(J)** Left: Example traces of EPSCs from vehicle-injected and lesioned animal in response to 5Hz stimulus train. Scale bar = 100pA∙125ms. Right: average EPSC amplitudes, normalized to the first EPSC, in response to 5Hz stimulation. (**K)** Left: Example traces of EPSCs from vehicle-injected and lesioned animal in response to 10Hz stimulus train. Scale bar = 100pA∙125ms. Right: average EPSC amplitudes, normalized to the first EPSC, in response to 10Hz stimulation. (Vehicle L2/3 neurons N=7, n=16, 6OHDA L2/3 neurons N=8 n=13; data shown as mean±SEM; *denotes p<0.05).

To begin investigating the synaptic mechanisms underlying this reduction in Mthal-EPSC amplitude, we calculated the coefficient of variation (1/CV^2^) and variance-to-mean ratio (VMR) in each group. The coefficient of variation from lesioned animals was reduced compared to controls (**Fig. 2F**, 1/CV^2^: Vehicle = 356.94±98.01, 6OHDA = 71.04±16.53, p=0.015), while there was no significant difference in VMR (**Fig. 2G**, VMR: Vehicle = 2.86±0.56, 6OHDA = 3.78±0.76, p=0.33). Previous studies interpreted a shift in 1/CV^2^, with no change in VMR, as a shift in presynaptic mechanisms, possibly a change in release sites (van Huijstee and Kessels, 2020). We next compared the input-output relationship at this synapse by varying stimulation intensity. Mthal-EPSCs onto L2/3 neurons were reduced at all intensities in lesioned animals compared to their control counterparts. This effect was present whether data were compared across individual cells or across animals within each experimental group, (**Fig. 2H-I**).

To further assess the consequence of dopamine depletion on Mthal transmission, we delivered trains of light stimuli at 5Hz and 10Hz frequency (**Fig 2J-K**) and quantified the short term dynamics of Mthal-EPSCs. Consistent with the driver role of Mthal input (Sherman, 2007), evoked EPSCs showed short term depression in both vehicle and lesioned mice. While there was no significant difference in short-term depression between groups in response to 5Hz stimulation, 10Hz trains of stimuli unveiled a significantly larger synaptic depression in the second and third EPSC in lesioned animals (**Fig 2K**, paired-pulse ratio of EPSC_2_/EPSC_1_: Vehicle = 0.71±0.038, 6OHDA = 0.57±0.048, p=0.037). These data support the interpretation that midbrain dopamine depletion alters synaptic transmission at the Mthal-M1 input in L2/3 and further support the interpretation that these changes are expressed presynaptically. Taken together, our findings demonstrate reduced baseline transmission and frequency-dependent impairments of short-term dynamics of Mthal-M1 transmission following midbrain dopamine depletion.

### Dopamine depletion impacts Mthal synaptic short-term depression in L5

To assess whether the effect of midbrain dopamine depletion on the Mthal-M1 connection may be layer-specific, we performed whole-cell recordings of L5 neurons and optically stimulated Mthal terminal fields (**Fig. 3**). L5 neurons included in the analysis showed regular firing pattern, pyramidal morphology, and lacked GAD67 expression (**Fig. 3A-B**). In contrast to L2/3, there was no significant difference in the amplitude of baseline Mthal-EPSCs onto L5 pyramidal neurons (**Fig. 3C** and second data point in panel H, in pA: Vehicle = 450.82±64.65, 6OHDA = 433.70±55.00, p=0.85). Similarly, there was no difference in either EPSC charge (**Fig. 3D**, in pA∙ms: Vehicle = 9077.54±1566.94, 6OHDA = 7417.00±992.31, p=0.43), or decay τ of scaled responses (**Fig. 3E**, in ms/pA: Vehicle = 0.042±0.0071, 6OHDA = 0.033±0.010, p=0.46). In accordance with unaltered Mthal-L5 synaptic strength, EPSC coefficient of variation and VMR were comparable between groups (**Fig. 2F** 1/CV^2^: Vehicle = 265.11±77.24, 6OHDA = 146.44±25.32, p=0.22) (**Fig, 2G** VMR: Vehicle = 3.39±0.46, 6OHDA = 4.00±0.65, p=0.44). Furthermore, increasing LED intensity did not reveal any differences in Mthal EPSCs onto L5 cells between experimental groups, whether quantified across neurons or across animals (**Fig. 3H-I** respectively). These results indicate that in contrast to Mthal input onto L2/3 cells, the baseline Mthal input strength to L5 neurons is unchanged. The laminar-specific effects of midbrain dopamine depletion may disrupt balanced thalamocortical activation of superficial and deep layers, an effect that could have significant functional consequences for the output of each circuit and their downstream targets.

**Figure 3.**
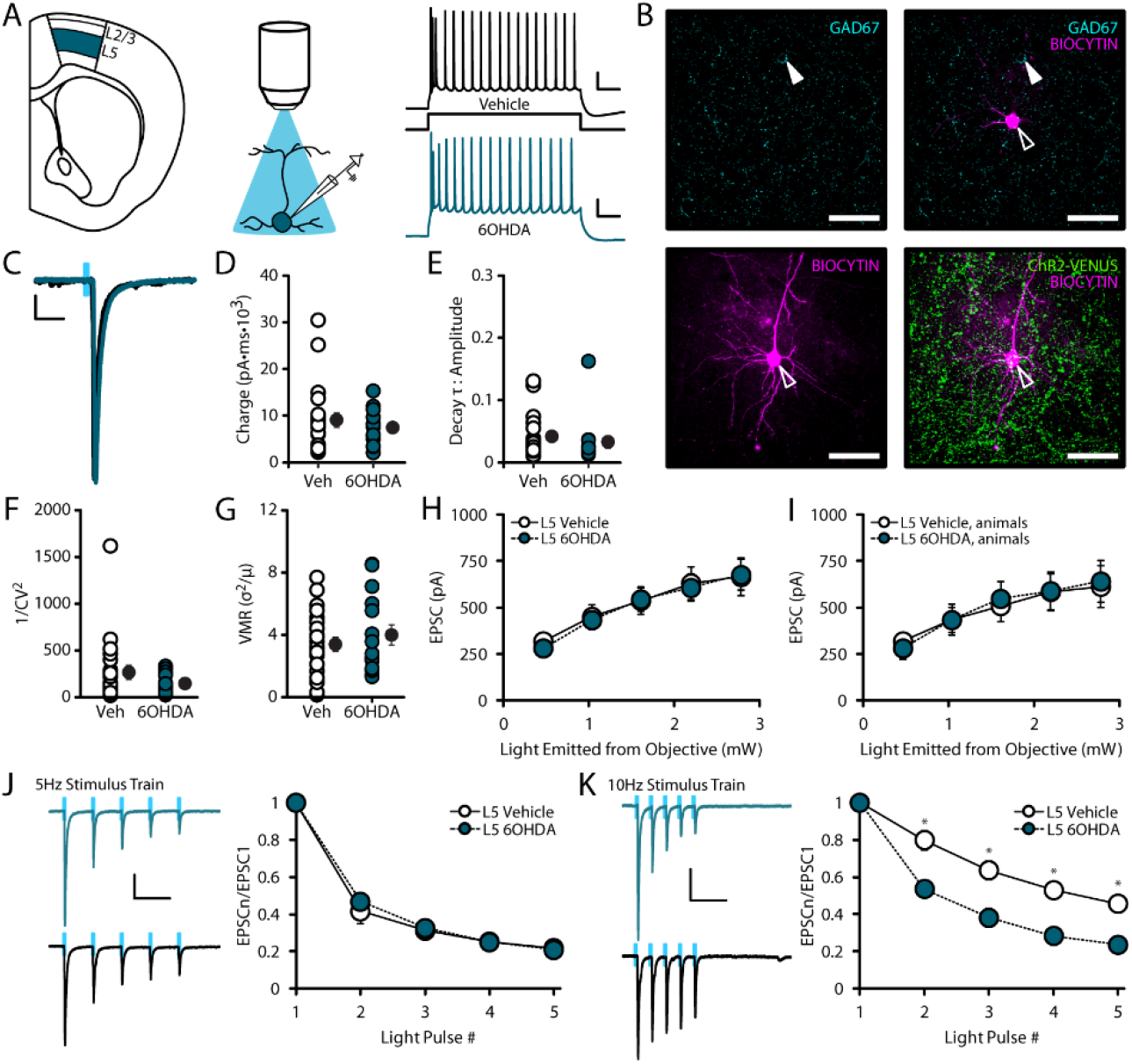
Mthal input to M1 L5 neurons is largely preserved in dopamine-depleted animals. **(A)** Whole-cell recordings of L5 pyramidal neurons in M1, with optical activation of ChR2-expressing thalamocortical axons. Right panel: typical firing pattern of a L5 neuron in vehicle-injected and lesioned animal. Scale bars = 20V∙100ms **(B)** Confocal images at 60x of an example recorded L5 neuron. Top panel: GAD67^−^recorded neuron (open arrow) with adjacent unrecorded GAD67^+^ neurons (filled arrows) and single z-plane depth. Bottom panel left: L5 recorded neuron shown as a collapsed stack spanning cell’s full structure. Bottom panel right: merge of L5 neuron and surrounding ChR2-expressing axons. Scale bars = 50μm. **(C)** Superimposed example traces of thalamocortical EPSCs from a vehicle-injected (black) and lesioned (teal) animal. Scale bar = 50pA∙50ms. **(D)** Calculated EPSC charges, and the group averages (black circles) using 5% LED intensity. **(E)** Calculated EPSC decay tau, normalized to EPSC amplitude, and the group averages (black circles) using 5% LED intensity. **(F)** Average coefficient of variation (1/CV^2^) of EPSCs in response to 5% LED intensity. **(G)** Average variance-to-mean raito (VMR) of EPSCs in response to 5% LED intensity. **(H-I)** Average input-output curves of EPSC amplitude in response to increasing LED intensity. **(J)** Left: Example traces of EPSCs from vehicle-injected and lesioned animal in response to 5Hz stimulus train. Scale bar = 100pA∙250ms. Right: average EPSC amplitudes, normalized to the first EPSC, in response to 5Hz stimulation. (**K)** Left: Example traces of EPSCs from vehicle-injected and lesioned animal in response to 10Hz stimulus train. Scale bar = 100pA∙250ms. Right: average EPSC amplitudes, normalized to the first EPSC, in response to 10Hz stimulation. (Vehicle L5 neurons N=9, n=21, 6OHDA L2/3 neurons N=6 n=15; data shown as mean±SEM; *denotes p<0.05).

Repetitive stimulation of thalamocortical inputs revealed a frequency-dependent change in short-term dynamics at these synapses. Short-term synaptic depression was comparable between experimental groups for trains of stimuli at 5Hz (**Fig. 3J**), while there was a significant increase in short-term depression in response to 10Hz stimulus trains in lesioned animals (**Fig 3K**, paired-pulse ratio of EPSC_2_/EPSC_1_: Vehicle = 0.80±0.052, 6OHDA = 0.54±0.047, p=0.0014). This effect suggests that while baseline Mthal drive onto L5 neurons are preserved following midbrain dopamine depletion, the activation of these synapses by patterned stimulation is impaired. The frequency-dependent increase in synaptic depression in the absence of change in CV suggests that this impairment engages distinct mechanisms from those we reported for L2/3.

### Mthal drive onto PV^+^ interneurons is preserved in dopamine-depleted animals

A core feature of thalamocortical drive is the engagement of cortical inhibitory interneurons. While thalamocortical afferents excite multiple types of inhibitory interneurons, in many cortices the largest thalamic drive is onto parvalbumin-expressing (PV^+^) cells (Swanson and Maffei, 2019). PV^+^ interneurons make perisomatic synapses onto glutamatergic neurons and are known for mediating powerful feedforward inhibition (Neske et al., 2015). We investigated how chronic midbrain dopaminergic cell loss impacts Mthal input onto M1 PV^+^ interneurons by performing whole-cell recordings in PVcretdtomato mice (**Fig. 4**). Recorded cells in L2/3 and L5 exhibited the fast-spiking firing phenotype typical of PV^+^ neurons (**Fig. 4A**) (Nassar et al., 2015). Tdtomato expression was preserved after immunohistochemical procedures, and all the neurons included in the analysis showed non-pyramidal morphology (**Fig. 4B**). Dopamine depletion had no effect on baseline Mthal-EPSCs onto L2/3 PV^+^ (**Fig. 4C, D**; second data point in D, in pA: Vehicle = 365.21±83.25, 6OHDA = 302.59±96.30, p=0.65) or L5 PV^+^ neurons (**Fig. 4G, H;** second data point in H, in pA: Vehicle = 389.25±84.81, 6OHDA = 261.65±48.48, p=0.19). Furthermore, we saw no effect of 6OHDA lesion on the input/output curve of Mthal-EPSCS onto L2/3 PV^+^ cells (**Fig. 4D**), or L5 PV^+^ cells (**Fig. 4H**). There were also no significant changes in coefficient of variation or VMR onto L2/3 (**Fig. 4E**, 1/CV^2^: Vehicle = 184.56±55.92, 6OHDA = 169.58±86.45, p=0.90) (**Fig. 4F**, VMR: Vehicle = 2.73±0.74, 6OHDA = 4.90±1.37, p=0.23), nor onto L5 PV^+^ neurons (**Fig. 4I**, 1/CV^2^: Vehicle = 128.80±40.69, 6OHDA = 77.01±19.10, p=0.24) (**Fig. 4J**, VMR: Vehicle = 5.97±2.10, 6OHDA = 4.40±0.95, p=0.47). Patterned Mthal terminal field stimulation at 5Hz and 10Hz evoked depressing EPSCs in both lesioned and control animals, however the degree of synaptic depression was not affected by a loss of midbrain dopamine (**Fig. 4K-L**). These results indicate that while Mthal input onto excitatory neurons is reduced following dopamine depletion, engagement of the PV^+^ inhibitory circuit is preserved.

**Figure 4.**
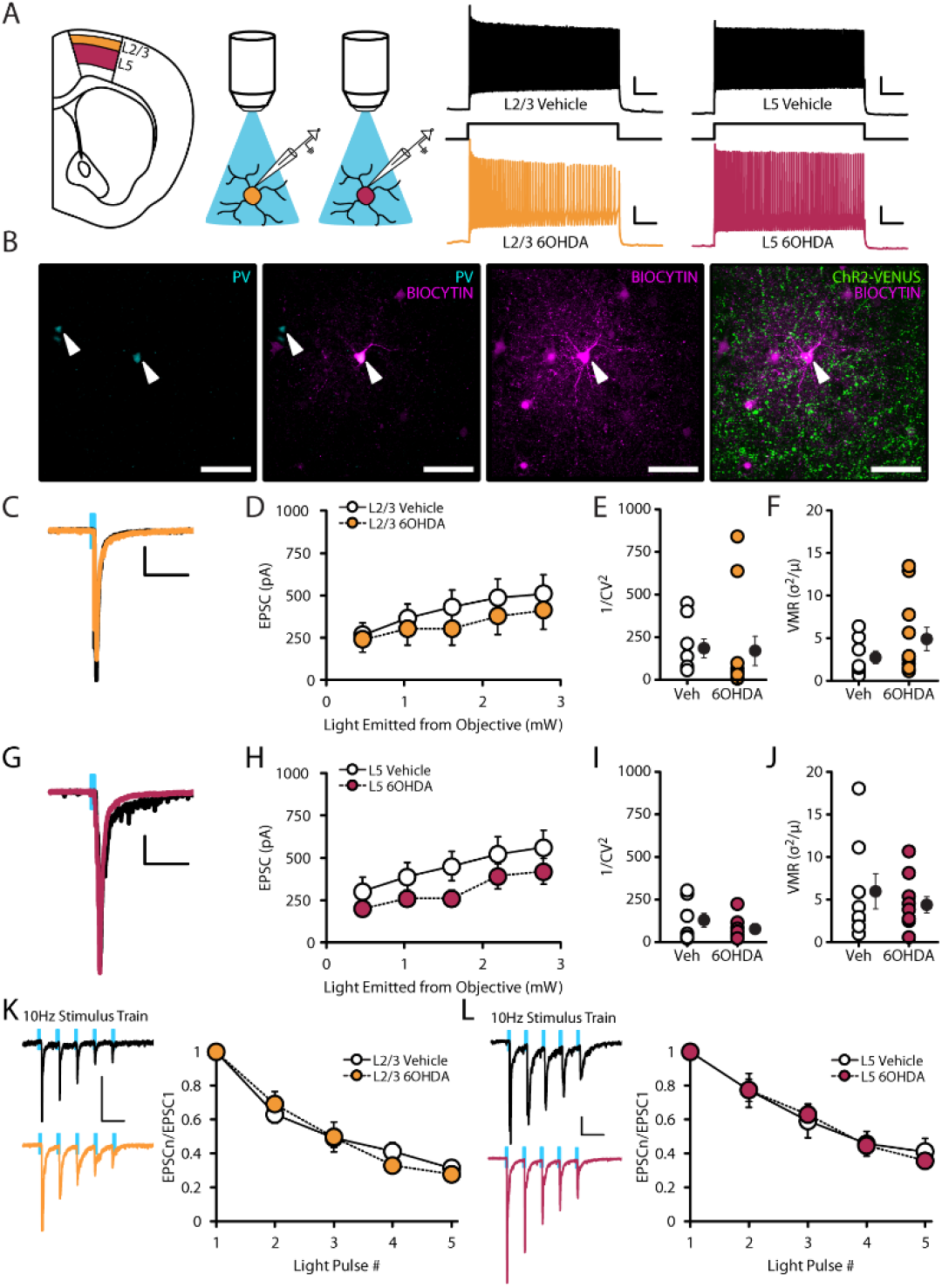
Mthal input on PV^+^ interneurons is not affected by dopamine depletion. **(A)** Whole-cell recordings of L2/3 and L5 PV^+^ neurons in M1, with optical activation of ChR2-expressing thalamocortical axons. Right panel: fast-spiking firing pattern of a L2/3 and L5 PV^+^ neurons in vehicle-injected and lesioned animals. Scale bars = 20mV∙100ms **(B)** Confocal images at 60x of an example recorded L5 PV^+^ neuron. Left panels: Tdtomato expression in the recorded neuron and adjacent unrecorded neuron (filled arrows), and merge image with biocytin labeling of recorded neuron GAD67^−^recorded neuron at a single z-plane depth. Right panel: L5 PV^+^ recorded neuron shown as a collapsed stack spanning cell’s full structure, and merged image with surrounding ChR2-expressing axons. Scale bars = 50μm. **(C)** Superimposed example traces of thalamocortical EPSCs in L2/3 PV^+^ neurons from a vehicle-injected (black) and lesioned (orange) animal. Scale bar = 50pA∙50ms. **(D)** Average input-output curves of evoked EPSC amplitudes in L2/3 PV^+^ neurons. **(E)** Average coefficient of variation (1/CV^2^) of EPSCs in response to 5% LED intensity. **(F)** Average variance-to-mean raito (VMR) of EPSCs in response to 5% LED intensity. **(G)** Superimposed example traces of thalamocortical EPSCs in L5 PV^+^ neurons from a vehicle-injected (black) and lesioned (magenta) animal. Scale bar = 50pA∙50ms. **(H)** Average input-output curves of evoked EPSC amplitudes in L5 PV^+^ neurons. **(I)** Average coefficient of variation (1/CV^2^) of EPSCs in response to 5% LED intensity. **(J)** Average variance-to-mean raito (VMR) of EPSCs in response to 5% LED intensity. **(K)** Left: superimposed example traces of L2/3 PV^+^ EPSCs from vehicle-injected and lesioned animal in response to 10Hz stimulus train. Scale bar = 50pA∙125ms. Right: average EPSC amplitudes, normalized to the first EPSC, in response to 10Hz stimulation. (**L)** Left: superimposed example traces of L5 PV^+^ EPSCs from vehicle-injected and lesioned animal in response to 10Hz stimulus train. Scale bar = 50pA∙125ms. Right: average EPSC amplitudes, normalized to the first EPSC, in response to 10Hz stimulation. (Vehicle L2/3 PV^+^ neurons N=4, n=8, 6OHDA L2/3 PV^+^ neurons N=8 n=11; Vehicle L5 PV^+^ neurons N=4, n=8, 6OHDA L5 PV^+^ neurons N=6 n=10; data shown as mean±SEM; *denotes p<0.05).

### M1 excitatory neuron output is enhanced in dopamine depleted animals

The magnitude of Mthal-EPSCs onto L2/3 excitatory neurons, and the short-term dynamics at Mthal-M1 synapses in both L2/3 and L5 are impacted by midbrain dopamine depletion. With reduced Mthal input to M1 excitatory neurons in mind, we investigated if these changes coincided with shifts in the input/output transformation of these cells (**Fig 5 A-D**: L2/3; **Fig 5E-H**: L5). We addressed this question by stimulating Mthal terminal fields at 5% LED intensity while recording excitatory postsynaptic potentials (Mthal-EPSPs) from L2/3 and L5 neurons, and comparing their magnitude to that of evoked currents at the same stimulation intensity (**Fig. 5A**: L2/3; **Fig. 5E**: L5). Both layers contained neurons with a variety of responses to Mthal stimulation: while some fired an action potential, others showed subthreshold EPSPs. First, we grouped L2/3 excitatory neurons with subthreshold Mthal responses and asked whether the magnitude of Mthal-EPSCs correlated with Mthal-EPSP amplitudes. In L2/3, there was a comparable correlation of Mthal-EPSCs and Mthal-EPSPs in control and 6OHDA mice (**Fig. 5B**; Vehicle R^2^ = 0.9169, 6OHDA R^2^ = 0.8063). Next, we quantified the proportion of L2/3 neurons that fired action potentials in response to Mthal stimulation in each experimental group (**Fig. 5C**) and found that Mthal axon stimulation evoked action potentials in a larger proportion of L2/3 pyramidal cells in 6OHDA animals relative to controls (**Fig. 5D**). These data suggest that despite reduced Mthal drive to L2/3 excitatory neurons, the transformation from synaptic input to output was enhanced. We observed a similar effect in L5: cells responding with subthreshold EPSPs showed a similar positive correlation between Mthal-EPSC and Mthal-EPSP amplitudes (**Fig. 5F**, Vehicle R^2^ = 0.2688, 6OHDA R^2^ = 0.4785). As observed in L2/3, a larger portion of L5 pyramidal neurons fired action potentials in response to Mthal stimulation in 6OHDA compared to vehicle-injected mice (**Fig. 5G, H**). These results suggest that dopamine depletion influences Mthal-evoked pyramidal neuron output in both superficial and deep layers of M1.

**Figure 5.**
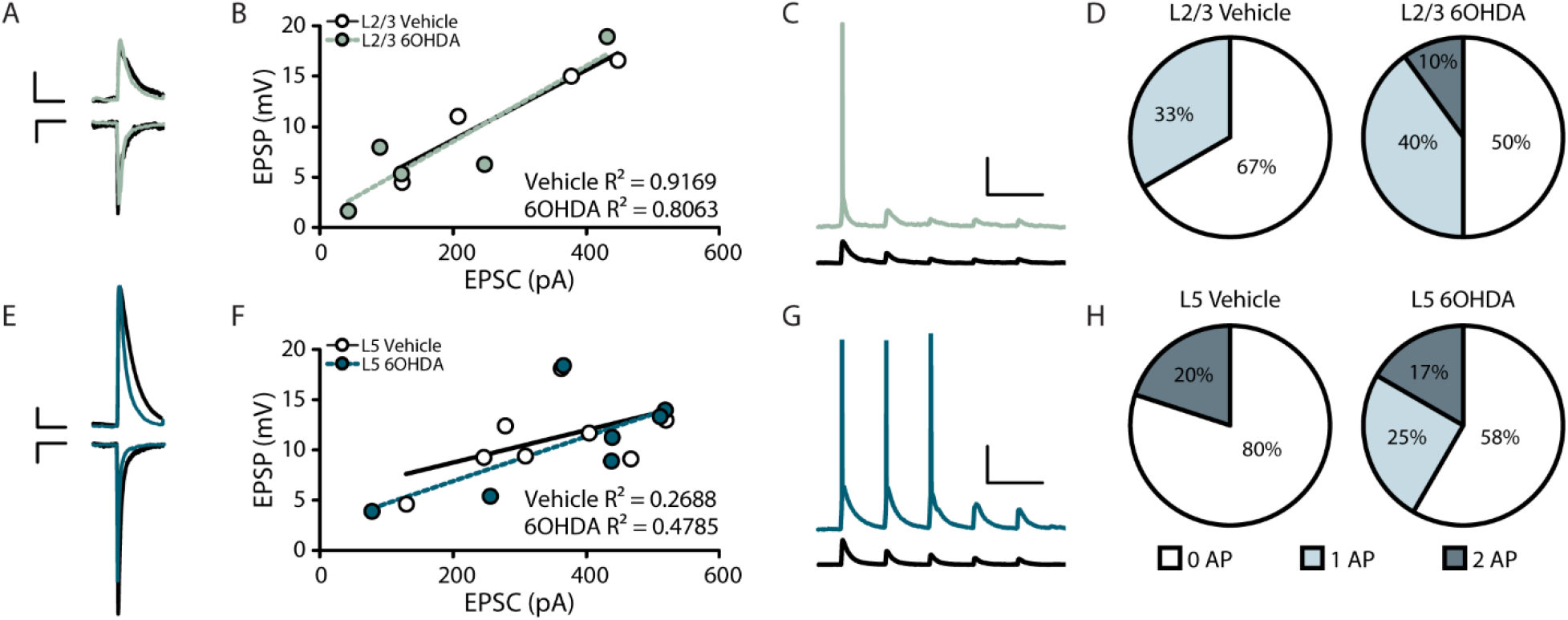
Midbrain dopamine depletion impacts M1 excitatory neuron output. **(A)** Mthal inputs were stimulated at 5% LED intensity and responses in L2/3 neurons were recorded in current clamp (EPSP) and voltage clamp (EPSC). EPSP Scale bar = 2.5mV∙125ms. EPSC Scale bar = 25pA∙125ms. **(B)** L2/3 EPSP amplitude and EPSC amplitude were positively correlated in both 6OHDA and control animals (Vehicle L2/3 neurons N=2 n=4, 6OHDA L2/3 neurons N=4 n=5). **(C)** 5Hz stimulation trains were delivered to stimulate Mthal axons and L2/3 responses were recorded in current clamp. Scale bar = 20mV∙250ms. **(D)** Pie charts showing the percentage of L2/3 neurons that firing 0, 1, or 2 action potentials (AP) in response to 5Hz Mthal stimulation. **(E)** L5 responses in current clamp (EPSP) and voltage clamp (EPSC) in response to 5% LED intensity Mthal axon stimulation. EPSP Scale bar = 2.5mV∙125ms. EPSC Scale bar = 50pA∙125ms. **(F)** L5 EPSP amplitude and EPSC amplitude were positively correlated in both 6OHDA and control animals (Vehicle L5 neurons N=5 n=8, 6OHDA L5 neurons N=4 n=7). **(G)** Current clamp responses of L5 neurons to 5Hz Mthal axon stimulation. Scale bar = 20mV∙250ms **(H)** Pie charts showing the percentage of L5 neurons firing in response to 5Hz Mthal stimulation.

The increase in the proportion of neurons firing action potentials in response to Mthal stimulation may be due to a selective shift in the excitability of these neurons. To test this possibility, we examined the input resistance and firing frequency of L2/3 and L5 neurons (**Supplementary Fig. 1**) included in the analysis of Mthal-EPSPs. There was no significant difference in input resistance, nor of frequency-current (*fI*) curves in cells from either layer (**Supplementary Fig. 1A-F**). These data suggest that changes in intrinsic excitability of M1 pyramidal neurons does not explain the increase in the proportion of neurons firing in response to Mthal stimulation.

An alternative possibility is that midbrain dopamine depletion shifts the balance between excitation/inhibition (E/I balance) onto M1 pyramidal neurons. To investigate this, we recorded spontaneous synaptic excitatory (sEPSCs) and inhibitory (sIPSCs) currents onto L2/3 and L5 excitatory neurons (**Fig. 6**). By recording at the reversal potential for Cl^−^ (−50mV), and at the reversal potential for cations mediating excitatory currents (+10mV), we isolated sEPSCs and sIPSCs and measured the amplitude and frequency of these events (**Fig. 6A-B**). In L2/3, sEPSC amplitude was reduced in 6OHDA animals (**Fig. 6C1**, KStest p=1.19E-7), while there was no change in EPSC frequency (**Fig. 6C2**, KStest p=0.60). Inhibitory synaptic activity onto L2/3 neurons was diminished in both amplitude (**Fig. 6C3**, KStest p= 1.24E-21), and frequency (**Fig. 6C4**, KStest p= 0.039). These results suggest that a shift in E/I balance from reduced intracortical inhibition may drive the increased output of L2/3 neurons despite reduced Mthal input (**Fig. 2**). In L5, there was a significant increase in both magnitude and frequency of spontaneous excitatory events (**Fig. 6D1**, KStest p=8.11E-16; Fig. **6D2**, KStest p=0.00068), while there was no change in spontaneous inhibitory activity (**Fig. 6D3**, KStest p=0.13; Fig. **6D4**, KStest p=0.57). These data, taken together with the lack of changes in baseline Mthal EPSCs (**Fig. 3**), suggest that loss of midbrain dopamine increases L5 neuron output due to enhanced ongoing intracortical excitation. Thus, midbrain dopamine depletion has profound effects on synaptic transmission within M1. It induces laminar-specific changes in Mthal synaptic drive, specifically at the inputs to pyramidal neurons. Furthermore, it shifts intracortical activity within M1 in a manner that alters the E/I balance onto L2/3 and L5 neurons in a layer-specific fashion. Taken together, dopamine depletion impacts the recruitment of M1 by Mthal and shapes the cortical transformation of input to output.

**Figure 6.**
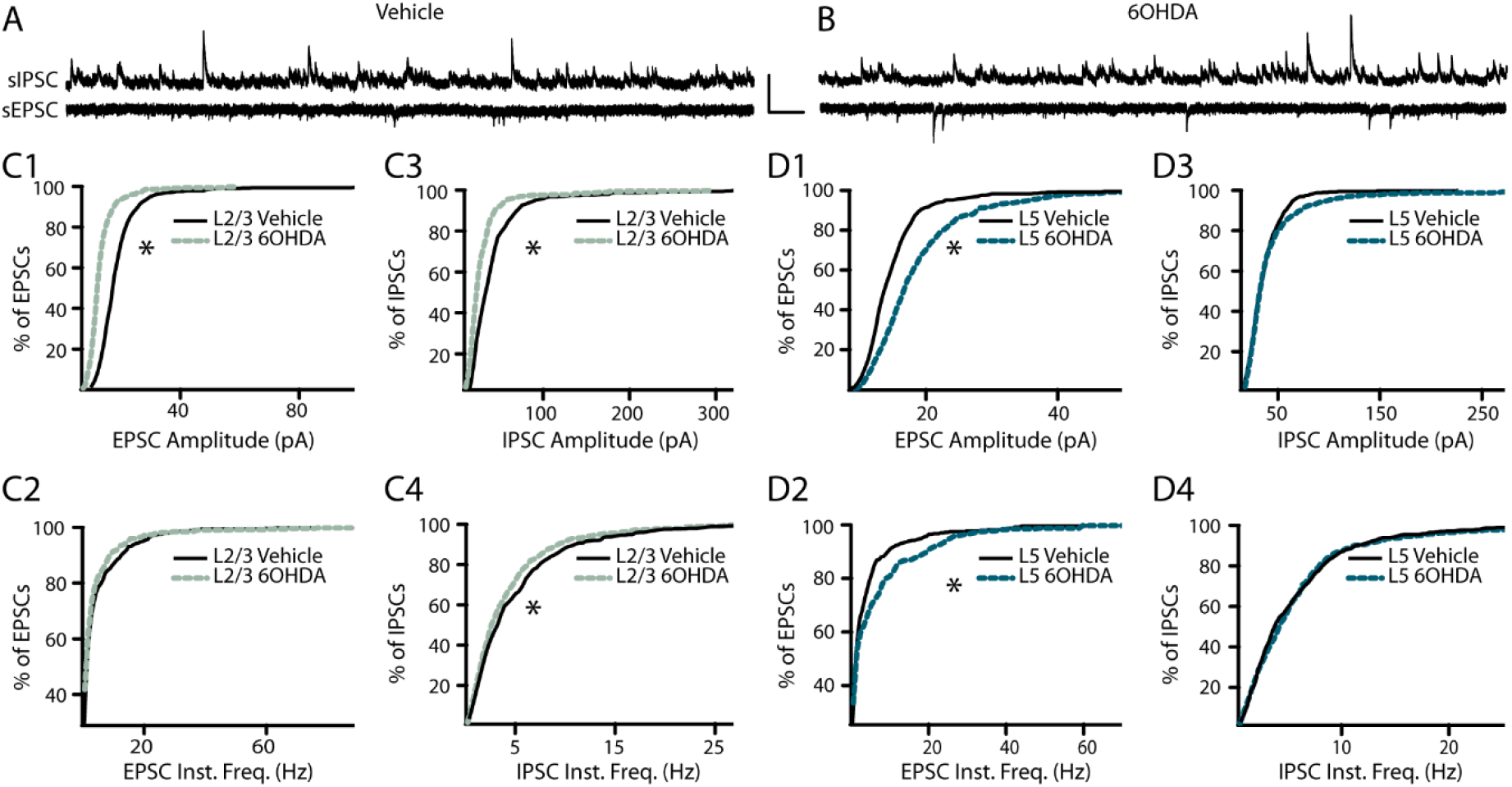
6OHDA lesions shifts the E/I balance onto M1 pyramidal neurons. **(A-B)** Example traces of isolated spontaneous excitatory and inhibitory postsynaptic currents (sEPSC, sIPSC) recorded in control and 6OHDA animals. Scale bar = 100pA∙500ms. **(C)** Cumulative probability plots of L2/3 sEPSC amplitude **(C1)**, sEPSC instantaneous frequency **(C2)**, sIPSC amplitude **(C3)**, and sIPSC instantaneous frequency **(C4)**. **(D)** Cumulative probability plots of L5 sEPSC amplitude **(D1)**, sEPSC instantaneous frequency **(D2)**, sIPSC amplitude **(D3)**, and sIPSC instantaneous frequency **(D4)**. (Vehicle L2/3 neurons N=3 n=5, 6OHDA L2/3 neurons N=4 n=5, Vehicle L5 neurons N=4 n=4, 6OHDA L5 neurons N=2 n=4. * denotes p<0.05).

## Discussion

Mthal (Planetta et al., 2013; Du et al., 2018) and M1 (Lefaucheur, 2005; Calabresi and Di Filippo, 2015; Guo et al., 2015) have been previously implicated as sites of dysfunction in PD. The accepted view of PD pathophysiology predicts that dopaminergic cell loss triggers widespread shifts in synaptic transmission along the motor pathway, resulting in abnormal thalamocortical excitation (Albin et al., 1989; Blandini et al., 2000; Barroso-Chinea et al., 2008; Braak and Del Tredici, 2008). However, the impact of dopamine loss on the synaptic physiology of the Mthal-M1 pathway had not been directly tested. Our findings corroborate these predictions and provide the first synaptic and circuit mechanisms demonstrating how cell loss within midbrain dopamine centers influences thalamocortical activation of M1 and its output.

### Layer-Specific Reduction in Mthal-M1 drive

6OHDA lesions of the SNc diminish Mthal drive onto excitatory L2/3, but not L5, excitatory neurons, indicating that the impact in M1 is layer-specific. This distinction is important when considering the differential role of neurons in L2/3 versus L5 of M1. *In vivo* imaging studies show that while both L2/3 and L5 neurons are active during movement, the activity of L2/3 excitatory cells is modulated by shifts in sensory experience, while L5 neuron activity correlated with the magnitude of motor response (Huber et al., 2012; Masamizu et al., 2014; Heindorf et al., 2018). Along with evidence that L2/3 receives major input from sensory cortices (Hooks et al., 2013), while L5 contains a high density of pyramidal tract neurons (Oswald et al., 2013; Jara et al., 2014), this suggests that L2/3 neurons are wired to integrate sensory cures into the motor response, while L5 activity underlies the “output” function of M1. Our data indicates that reduced Mthal drive to L2/3 in 6OHDA animals may impair sensorimotor integration in M1, while Mthal drive within the output layer is preserved.

Analysis of Mthal-evoked EPSC amplitudes in L2/3 revealed a reduction in the coefficient of variation with no change in VMR. These parameters provide insights into the site of expression and possible mechanisms underlying a change in synaptic strength (Brock et al., 2020). A previous study reported that decreased EPSC amplitude, coupled with decreased coefficient of variation and unchanged VMR indicates that reduction of release sites is the likely mechanism (van Huijstee and Kessels, 2020). According to this analysis, the reduction in Mthal-EPSC amplitude in L2/3 following midbrain dopamine depletion depends on presynaptic changes.

### Loss of dopamine and Mthal-M1 synapse dynamics

Short-term plasticity regulates the response strength of cortical neurons to fast, patterned stimulation (Hennig, 2013). Synaptic depression is a form of short-term plasticity common to driver thalamocortical pathways (Sherman, 2007), which aids in cortical processing by decorrelating coinciding inputs to reliably follow patterned activity (Blackman et al., 2013). The magnitude of short-term depression of synaptic responses is influenced by changes in release probability, receptor desensitization, or calcium dynamics (Fioravante and Regehr, 2011). Our results show that 6OHDA lesions of the SNc lead to increased synaptic depression at Mthal synapses onto M1 pyramidal neurons in both L2/3 and L5. In L2/3, this phenomenon was accompanied by decreased 1/CV^2^, further supporting the interpretation that in this layer, the shift in short-term dynamics was due to a presynaptic change. In L5, the increased synaptic depression occurred independent of any changes to 1/CV^2^, suggesting that a different mechanism, likely postsynaptic, may be driving the increase in short-term depression at these synapses (Schneggenburger et al., 2002). Together, these results demonstrate that midbrain dopamine loss alters thalamocortical synaptic dynamics, an effect that is likely to profoundly affect M1 responsiveness to patterned stimuli.

### Mthal-M1 PV^+^ activation is preserved in PD model

Functionally, inhibitory signaling mediated by M1 PV^+^ interneurons is important for sculpting the tuning of adjacent pyramidal neurons (Merchant et al., 2008), the initiation of movements (Estebanez et al., 2017), and motor learning (Chen et al., 2015). Many of the functions of PV^+^ cells involve regulating pyramidal neuron output. We report that dopamine depletion did not alter drive or short-term plasticity at Mthal synapses onto PV^+^ interneurons. Reduced thalamocortical activation of excitatory neurons, coupled with preserved activation of neighboring PV^+^ interneurons, could have serious consequences on balanced synaptic transmission within M1. 6OHDA lesions have caused similar imbalances in other motor areas, including the basal ganglia. Like Mthal input to the cortex, glutamatergic cortical afferents in the striatum synapse onto both MSNs and PV^+^ cells (Ramanathan et al., 2002), the latter mediating powerful feedforward inhibition onto neighboring MSNs (Gittis et al., 2010). The function of this circuit is crucial for motor learning and reward-guided behaviors (Lee et al., 2017). In a 6OHDA model of PD, corticostriatal input to MSNs was affected by dopamine loss, while input to striatal fast-spiking interneurons (putatively PV^+^) was unchanged, worsening an imbalance between striatopallidal and striatonigral (indirect and direct pathway, respectively) neurons. This cell type-specific effect is comparable to the one we observed in M1. Preserved thalamocortical activation of PV^+^ neurons accompanied with altered feedforward excitation following dopamine depletion may impair M1 activation during voluntary movement and impair M1 output to its projection targets.

### Increased Mthal-evoked output of pyramidal neurons

Despite reduced Mthal drive to L2/3, we observed increased probability of Mthal-evoked firing in pyramidal neurons in both L2/3 and L5. Further analysis excluded changes in intrinsic excitability and pointed to layer-specific shifts in E/I balance as a possible mechanism underlying the increased output of M1 neurons. In L2/3, there was a decrease in amplitude and frequency of sIPSCs, suggesting that dopamine depletion has a net disinhibitory effect on these neurons. In contrast, in L5 there was an increase frequency and amplitude of sEPSCs. Both disinhibition and increased overall excitatory drive may explain the increased proportion of M1 neurons firing action potentials in response to Mthal stimuli. Thus, loss of midbrain dopamine results in impaired Mthal activation of M1 and shifts the E/I balance of intracortical activity. The combination of these changes results in altered input/output transformations in pyramidal neurons.

## Conclusion

The engagement of M1 neurons is crucial for many aspects of movement, including action observation (Vigneswaran et al., 2013), motor learning (Molina-Luna et al., 2009; Hosp et al., 2011; Kida et al., 2016), and skilled movement execution (Kaufman et al., 2013; Ueno et al., 2018). Studies focused on the circuit consequences of PD predicted that basal ganglia dysfunction leads to abnormal synaptic transmission along the Mthal-M1 pathway. Our findings that dopamine depletion shifts the drive and dynamics of Mthal-M1 synapses is a novel contribution to the involvement of M1 in PD pathophysiology. Further, we found that while thalamocortical engagement of PV^+^ interneurons is intact, recurrent inhibitory activity is altered, which opens new questions concerning the role of M1 inhibitory circuits during movement in a Parkinsonian state. Our findings indicate altered synaptic transmission, both long-range and recurrent, as a source of M1 dysfunction in an animal model of PD. This study corroborates existing models of PD and emphasizes how a loss of dopamine impacts the output center for voluntary movement.

## Acknowledgments

This work was supported by the Hartman Foundation for Parkinson’s disease.

---

**Supplementary Figure 1.**
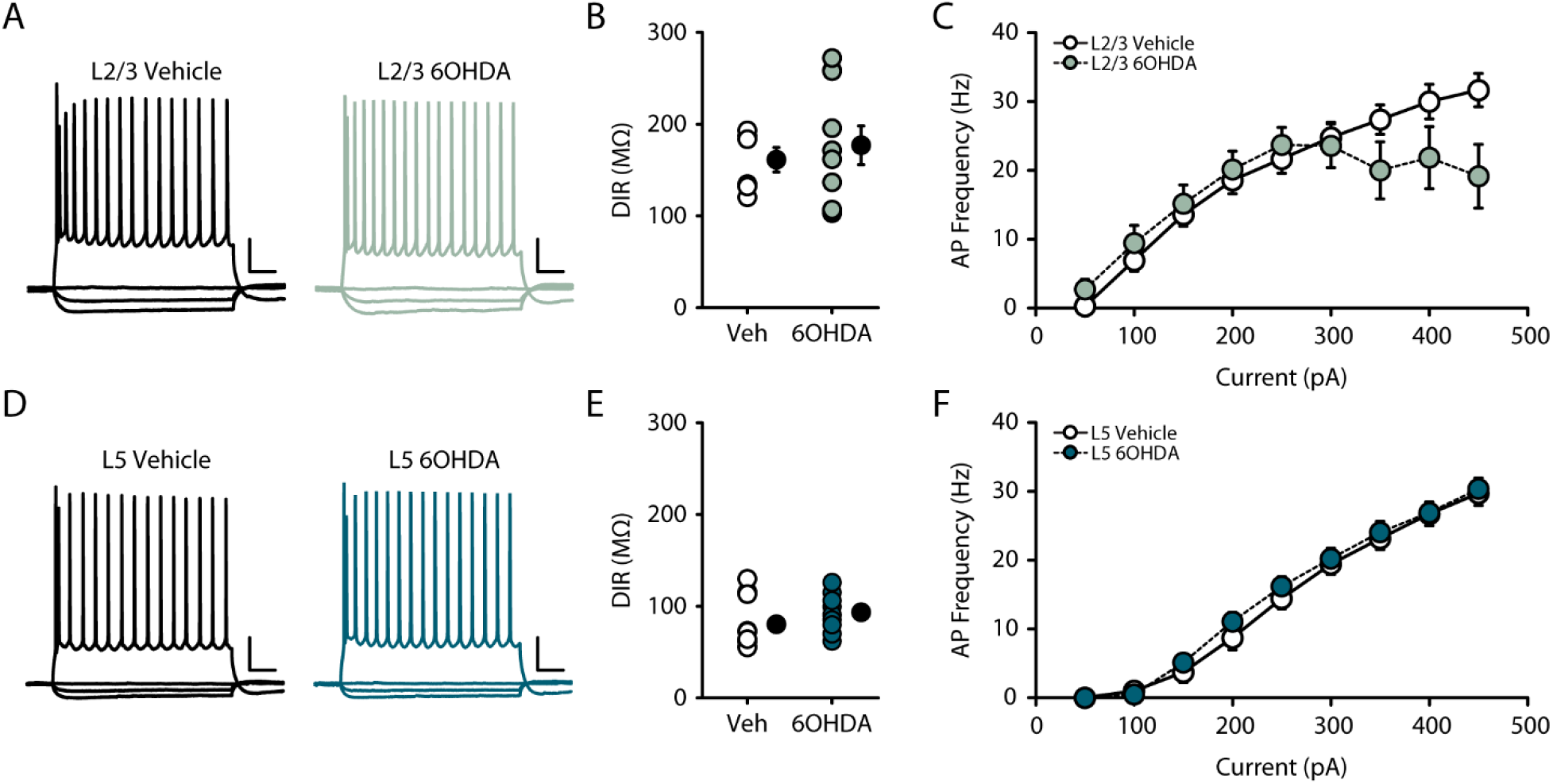
Pyramidal neuron excitability following 6OHDA lesion. **(A,D)** Example traces of L2/3 and L5 pyramidal neuron responses to hyperpolarizing and depolarizing current injection. (−100pA, −50pA, 0pA, and +350pA steps shown). Scale Bar = 20mV∙100ms. **(B,E)** Average dynamic input resistance in L2/3 and L5 neurons between experimental groups. **(C,F)** Average frequency-current curves for L2/3 and L5 neurons between experimental groups. (Vehicle L2/3 neurons N=3, n=6, 6OHDA L2/3 neurons N=8, n=10; Vehicle L5 neurons N=5, n=10, 6OHDA L5 neurons N=7, n=12; data shown as mean±SEM; *denotes p<0.05).

